# Mammalian cells evacuate and shelter mitochondrial DNA from destruction following hypoxia response-induced mitophagy

**DOI:** 10.1101/2025.08.01.667885

**Authors:** Reiko Kamegai, Asato Uenosono, Fumiyo Ikeda, Keisuke Ohta, Kota Yanagitani

## Abstract

Cells use a specialized process called mitophagy to degrade mitochondria, and the conditions that induce mitophagy can endanger mitochondrial DNA (mtDNA). Here, we report a novel mechanism by which cells undergoing hypoxia response (HR)-induced mitophagy protect their mtDNA from destruction by segregating it into large mitochondria (Mito-L). Using time-lapse imaging, we found that Mito-L are formed when mitochondria undergo tubular-to-spherical transitions and then fuse together. Under HR-activating conditions, we found the mitophagy process itself is essential for Mito-L formation. When we fragmented mitochondria prior to inducing the HR, we observed fewer Mito-L and more mtDNA degradation. Thus, mammalian cells protect mtDNA against excessive mitophagy induced by HR, thereby preventing the loss of the genetic information.

Hypoxia response-induced mitophagy triggers cells to shelter mitochondrial DNA (70 characters)

## Main text

Mitochondrial DNA (mtDNA) encodes essential subunits of the respiratory chain complexes required for oxidative phosphorylation (*1, 2*). This means loss of mtDNA prevents eukaryotic cells from producing ATP via oxidative phosphorylation. Modern eukaryotes seem to have inherited their mtDNA from an ancient ancestral eukaryote, suggesting there has been sufficient time and pressure to support the evolution of elaborate mechanisms for preventing mtDNA loss under harsh conditions. We therefore hypothesized the existence of a protective mechanism that safeguards mtDNA under conditions that induce mitophagy—the major cellular pathway for reducing mitochondria (*3-5*). To identify such a mechanism, we investigated mtDNA under hypoxic conditions that induce mitophagy as part of the hypoxia response (HR) (*6, 7*).

### The hypoxic response induces the production of large, spherical, mtDNA-enriched mitochondria (Mito-L) and small, spherical, mtDNA-free mitochondria (Mito-S)

Under hypoxic conditions, the limited availability of oxygen disrupts the respiratory chain, increasing the production of reactive oxygen species (ROS). To mitigate this, the HR includes mitophagy to shut down the respiratory chain (*8, 9*). When we examined mtDNA under conditions inducing the HR, we observed a segregation of mitochondria into two populations: large, spherical mitochondria with diameters > 1 μm and small, spherical mitochondria with diameters < 0.5 μm (Figs. 1A, S1A). The mitochondria of non-stressed HeLa cells typically exhibit a tubular morphology with discrete mtDNA. Under conditions inducing the HR, however, mtDNA staining seems to increase in the large, spherical mitochondria, leaving the small, spherical mitochondria nearly devoid of it. From now on, we will refer to these large, spherical mitochondria containing at least two mtDNA copies as Mito-L (Fig. S1B) and to the small, spherical mitochondria lacking mtDNA as Mito-S. To quantify Mito-L formation, we calculated the percentage of cells harboring Mito-L relative to the total cell population (Fig. 1A; Fig. S1C). HR is a cellular response activated by the HIF1 transcription factor. Under normoxic conditions, however, HR is not induced because Hypoxia-Inducible Factor 1α (HIF1α), a subunit of HIF, is rapidly degraded. In contrast, under hypoxic conditions, the degradation of HIF1α is inhibited, allowing it to form a heterodimer with the other subunit, Hypoxia-Inducible Factor 1β (HIF1β), thereby inducing HR (*10, 11*). Notably, HR activation can also be triggered under normoxic conditions by pharmacological stabilization of HIF1α using dimethyloxalylglycine (DMOG; *11*). We found that stabilization of HIF1α with DMOG induces a similar mitochondrial bipolarization in HeLa cells and human fibroblasts (L23immo; Fig. S1D) and that siRNA-mediated knockdown of HIF1α abolished DMOG-induced Mito-L formation (Fig. S1E), confirming a role for HIF1α-regulated transcription in this process.

**Fig. 1.**
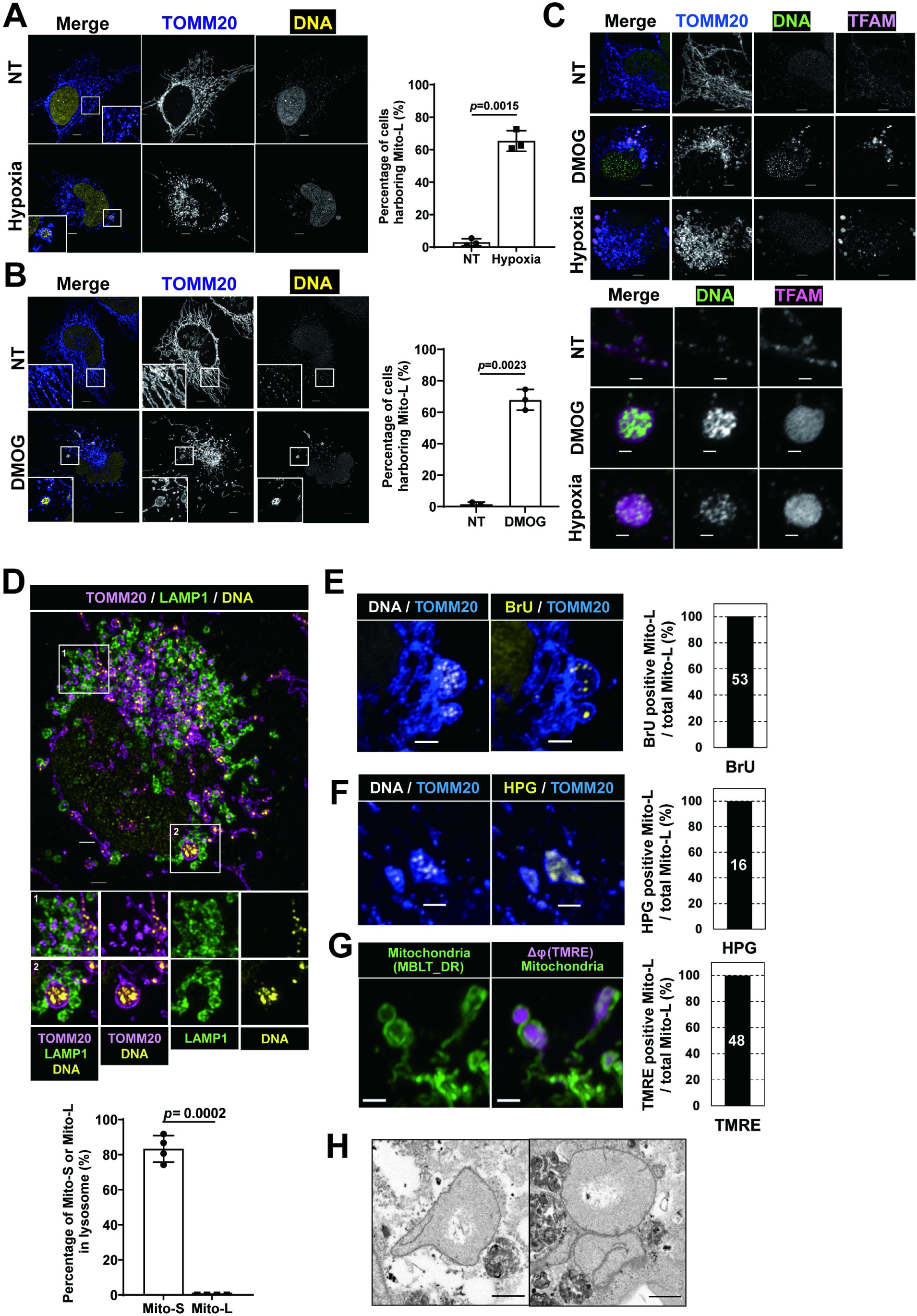
The hypoxic response leads to large, spherical, mtDNA-enriched mitochondria (Mito-L) and small, spherical, mtDNA-free mitochondria (Mito-S). (**A**) HeLa cells were incubated in normal or hypoxic DMEM for 24 h. The mitochondrial outer membrane and DNA were visualized via immunostaining with antibodies against TOMM20 and DNA, respectively. The right graph shows the percentage of cells harboring Mito-L from three independent experiments. (**B**) HeLa cells were treated with DMOG (1 mM) for 30 h. Immunostaining and graph generation were performed as in (A). (**C**) HeLa cells treated with the indicated stimuli as in (A and B) were stained with the indicated antibodies. (**D**) HeLa cells were treated with DMOG (1 mM) for 30 h. Immunostaining was performed with the indicated antibodies as in (A). The bottom graph shows the percentage of Mito-L or Mito-S within lysosomes. The data were collected from four independent cells. (**E-F**) BrU or HPG were incorporated into DMOG-treated cells and visualized via an anti-BrdU antibody or click-chemistry-based conjugation of Alexa Fluor 594, respectively. Those signals were then co-immunostained with the indicated antibodies. The right graphs indicate the percentage of Mito-L containing BrU (E) or HPG (F). (**G**) Mitochondria and the electrostatic membrane potential of the mitochondrial inner membrane in DMOG-treated HeLa cells were visualized with MitoBright LT Deep Red and TMRE, respectively. The right graph indicates the percentage of TMRE-positive Mito-L out of the total number of Mito-L. In panels A and B, the data are presented as means ± SD from three independent samples. Statistical analysis was performed with two-tailed Student’s t-tests. P values are indicated in each graph. (**H**) Electron micrographs of Mito-L in a HeLa cell treated with DMOG (1 mM) for 30 h. Accumulation of mtDNAs within the swollen mitochondria was confirmed by correlative light and electron microscopy (CLEM; Fig. S3A-B). Scale bars, 5 μm (A, B, C top), 2 μm (D-G), and 1 μm (C bottom, H).

Next, we found that it is mitophagy that leads to the segregation of these two mitochondrial subtypes. When we stained for mitochondrial and lysosomal markers, we found that Mito-S are predominantly (∼80%) sequestered in lysosomes (Fig. 1D). This indicates Mito-S are degradation intermediates. In contrast, Mito-L produced under DMOG treatment were not associated with lysosomes. Since Mito-L are enriched with mtDNA, we further investigated their nature by analyzing their gene expression profiles. Specifically, we evaluated their transcriptional and translational activity under DMOG-treated conditions by measuring their incorporation of 5-bromouridine (BrU) and L-homopropargylglycine (HPG), respectively. In the HPG incorporation experiment, we added cycloheximide to inhibit cytosolic translation and ensure exclusive labeling of mitochondrial translation. As shown in Figs. 1E-F and Figs. S1F-G, the Mito-L of DMOG-treated cells exhibited active transcription and translation. Furthermore, although the cristae membranes of Mito-L were more loosely packed (Figs. 1H, S1H, S2A-B and Movie S1), they exhibited normal polarization, as detected by TMRE staining (Fig. 1G). In the three-dimensional reconstructed model of Mito-L, the cristae membranes were displaced toward the periphery, and mtDNA was accumulated in the cristae-free space (Movie S1). In summary, we discovered that the HIF1-mediated gene expression program segregates tubular mitochondria into two distinct types of structures: gene-expression active Mito-L and degradative Mito-S.

### Dynamics of Mito-L formation

To better understand how Mito-L is formed, we performed time-lapse imaging of DMOG-treated HeLa cells. For this experiment, we labeled mitochondria and mitochondrial nucleoids comprising mtDNA and mtDNA binding proteins with EGFP-MAOA and TFAM-mScarlet, fluorescent proteins that label the mitochondrial outer membrane (MOM) and mitochondrial nucleoids, respectively (*12, 13*). Figure 2A shows how the normally tubular mitochondria passed through a tubular-to-spherical transition, producing intermediate-sized Mito-L (diameter: ∼1 μm; Movie S2). The resulting structures then fused with one another, producing even larger Mito-L (diameter: ∼2 μm; Movie S3; Fig. S3A).

**Fig. 2.**
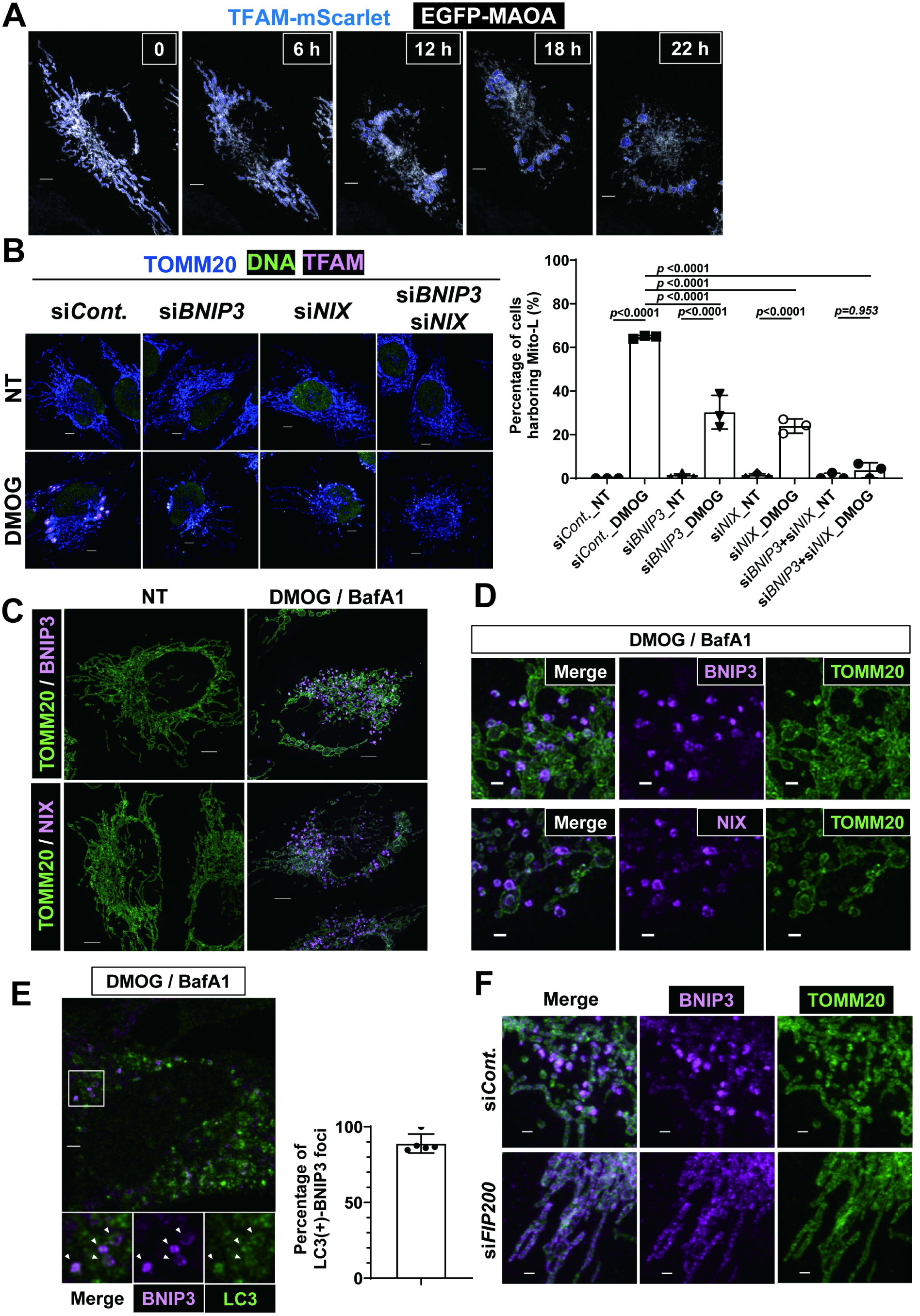
Mito-L formation depends on BNIP3/NIX-mediated mitophagy. (**A**) Time-lapse imaging of HeLa cells transiently expressing TFAM-mScarlet and EGFP-MAOA in the presence of DMOG (1 mM). (**B**) HeLa cells treated with the indicated siRNAs were subjected to DMOG (1 mM) treatment for 30 h. Mito-L formation was visualized via immunostaining with the indicated antibodies. The right graph shows the percentage of cells harboring Mito-L. The data are presented as means ± SD from three independent samples. Significant differences were calculated using two-way ANOVA with Tukey’s tests for multiple comparisons. (**C, D**) HeLa cells were treated with DMOG (1 mM) and Bafilomycin A1 (BafA1; 100 nM) for 12 h. Immunostaining was performed with the indicated antibodies. (**E**) HeLa cells were treated with DMOG and BafA1 as in (C, D). Incorporation of BNIP3 vesicles into autophagosomes was assessed via immunostaining with the indicated antibodies. Arrowheads indicate BNIP3-positive vesicles within LC3 vesicles. The right graph shows the percentage of BNIP3-positive vesicles within LC3 vesicles out of the total number of BNIP3-positive vesicles. These data were collected from five different cells. (**F**) HeLa cells treated with the indicated siRNAs were subjected to treatment with DMOG (1 mM) and BafA1 (100 nM) for 12 h. BNIP3 distribution was visualized via immunostaining with the indicated antibodies. Scale bars, 5 μm (A, B, C top), 2 μm (D-G), and 1 μm (C bottom).

### BNIP3- or NIX-mediated mitophagy initiates Mito-L formation during HR activation

So far, we have shown that Mito-L formation requires the hypoxic response (HR) mediated by HIF1; however, whether the induction of mitophagy is a direct cause has not been determined. Therefore, we investigated the roles of BNIP3 and NIX/BNIP3L—major mitophagy receptors and transcriptional targets of HIF1—in Mito-L formation (*9, 14, 15*). Individual knockdown of either BNIP3 or NIX reduced Mito-L formation under DMOG treatment, but simultaneous knockdown of both BNIP3 and NIX almost completely abolished Mito-L formation (Fig. 2B, S3B). We also found that FIP200, a critical autophagy factor, was necessary for Mito-L formation during hypoxia (Fig. S3C, S3D). Thus, BNIP3- and NIX-mediated mitophagy is essential for Mito-L formation under hypoxic conditions.

To understand how mitophagy contributes to Mito-L formation, we examined the distribution of BNIP3 and NIX during DMOG-induced Mito-L formation. Using live-cell imaging, we found Mito-L began to form around 12 hours after the addition of DMOG (Fig. 2A, Movie S2). When we added Bafilomycin A1 in addition to DMOG to prevent mitophagosome degradation, we found increased levels of BNIP3 and NIX in the form of focal structures co-localizing with the mitochondrial marker TOMM20 (Fig. 2C, 2D, S3F). We confirmed the specificity of the BNIP3 and NIX signals by demonstrating their elimination with knockdown of each respective gene. Interestingly, we observed that the BNIP3 foci appeared both on tubular mitochondria and on pre-vesicular structures budding from the tubular mitochondria in a process that resembled secretory pathway vesicle formation (Fig. S3E). This raised the possibility that BNIP3 is initially localized on tubular mitochondria, subsequently recruited to the budding site, and then pinched off from the tubule along with the mitochondrial vesicles.

Given that BNIP3 is a mitophagy receptor and that almost all BNIP3 foci were present in autophagosomes, as visualized with an anti-LC3 antibody (Fig. 2E), we hypothesized that autophagy may be involved in the formation of BNIP3 foci. To test this, we knocked down the essential autophagy gene FIP200 in HeLa cells and then examined the resulting distribution of BNIP3. In these FIP200-deficient cells, we observed even distribution of BNIP3 across the entire mitochondrial structure, rather than the focal staining we observed in the presence of DMOG (Fig. 2F, S3G-H). Notably, the DMOG-induced increase in BNIP3 in FIP200-deficient cells was similar to that of control siRNA-treated cells (Fig. S3D).

We next analyzed the constituent proteins of BNIP3 vesicles. As shown in Fig. 3A, we found BNIP3 vesicles contained markers of the mitochondrial outer membrane (MOM), such as TOMM20 and TOMM40, as well as markers of the mitochondrial inner membrane (MIM), such as TIMM23 and ATP5A. BNIP3 vesicles also contained modest amounts of mitochondrial matrix components, such as HSP60 and citrate synthase. Interestingly, however, we found that mt-nucleoid components like mtDNA and TFAM were absent from most of the BNIP3 foci (Fig. 3A, B). This suggests nucleoids tend to avoid incorporation into BNIP3 vesicles.

**Fig. 3.**
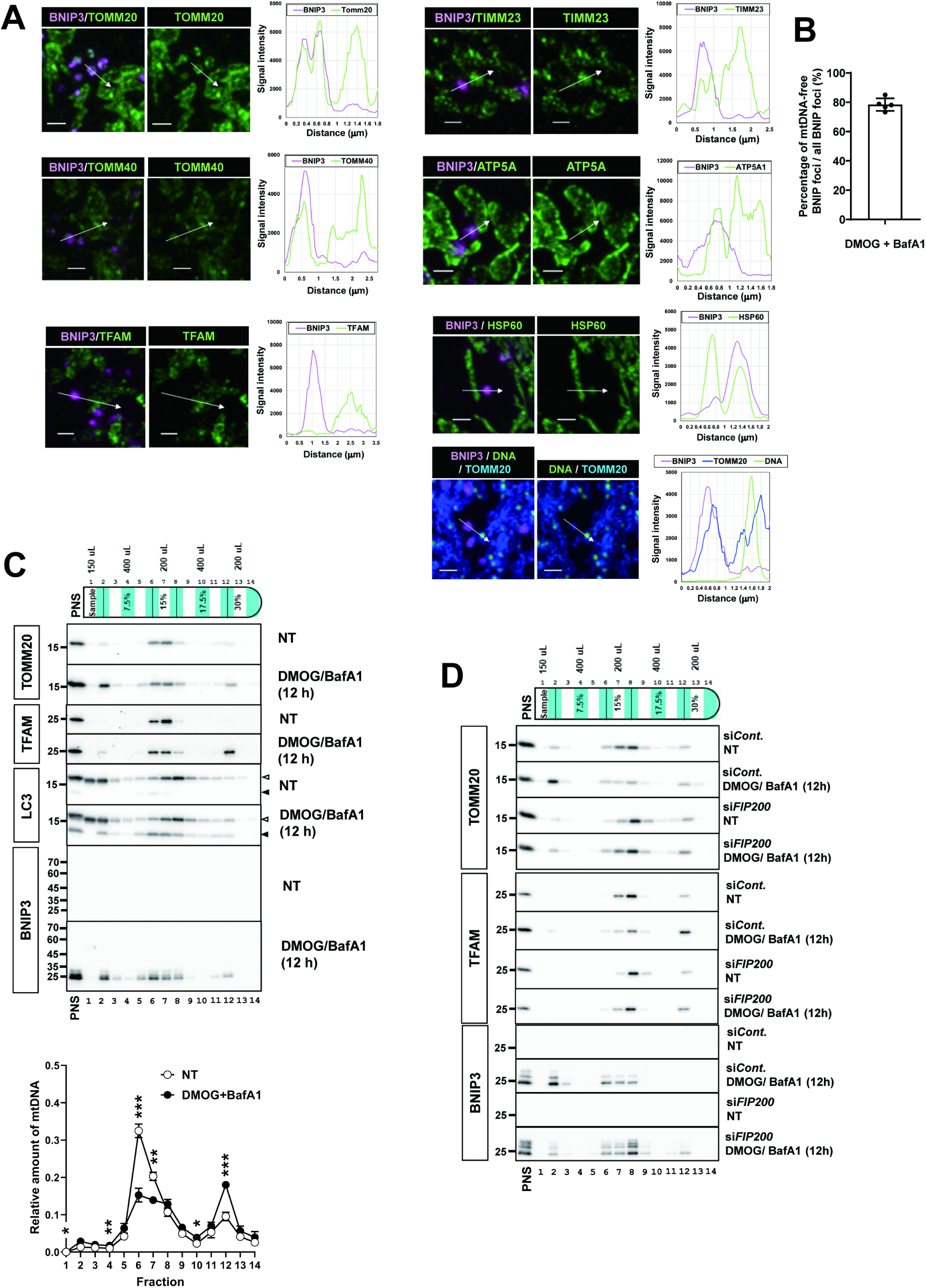
The majority of BNIP3-positive mitophagosomes do not contain mtDNA. (**A**) HeLa cells transiently expressing TFAM-mScarlet and EGFP-MAOA were treated with DMOG and BafA1 as in Fig. 2C and immunostained with the indicated antibodies. The right panels show the fluorescence intensity of each channel plotted along the length of the white arrow. Time-lapse imaging of those signals was carried out in the presence of DMOG (1 mM). (**B**) The percentage of DNA-free BNIP3 foci among all BNIP3 foci was calculated from five independent cells. (**C**) HeLa cells were treated with or without DMOG (1 mM) and BafA1 (100 nM) for 12 h. Cell homogenates were then separated by iodixanol density gradient centrifugation. The distribution of the indicated proteins (top panels) or mtDNA (bottom graph) were analyzed by western blot using the corresponding antibodies or by quantitative PCR with the corresponding primer sets. The bottom graph indicates the relative distribution of mtDNA per fraction as assessed by quantitative PCR in four independent experiments. For each fraction, significant differences between paired groups were calculated using Student’s two-tailed unpaired t-tests. P values are indicated (\**P* < 0.01, \*\**P* < 0.001, \*\**P* < 0.0001). (**D**) HeLa cells treated with the indicated siRNAs were subjected to DMOG (1 mM) and BafA1 (100 nM) treatment for 12 h. Cell homogenates were separated and analyzed as in (C). Scale bars, 1 μm (A, B).

Together, we propose a model for BNIP3 vesicle formation (Fig. S4A). Initially, BNIP3 is synthesized in the cytosol and transported to the MOM under HR-inducing conditions. During mitophagy, the ULK1 complex, including the FIP200 subunit, initiates the formation of a phagophore, whose surface is decorated with lipidated LC3. This then collects BNIP3 from tubular mitochondria via BNIP3’s affinity for LC3, forming a budding structure on the tube. The closing autophagosome then pinches off a BNIP3-enriched portion, forming BNIP3 vesicles destined for autolysosomal degradation.

Given that BNIP3 vesicles preferentially contained MOM and MIM markers while excluding mtDNA, their formation should enrich mtDNA in the remaining tubular mitochondria. We noted that TFAM levels were also higher in these remaining tubular mitochondria. Recent studies indicated that TFAM and mtDNA can undergo liquid-liquid phase separation (LLPS) when concentrated together (*16, 17*). It is therefore likely that the resulting increased density of mtDNA and TFAM in the remaining tubular mitochondria induces the formation of large spherical LLPS structures that eventually lead to the formation of Mito-L.

### Biochemical characterization of Mito-L and BNIP3 vesicles indicates that mtDNA is preferentially partitioned into Mito-L over BNIP3 vesicles

In our microscopic observations, Mito-Ls are large, spherical mitochondria rich in proteins (especially TFAM) and mtDNA. Given that DNA and proteins are high-density materials, Mito-Ls are presumably heavier than normal mitochondria. BNIP3 vesicles, in contrast, are small mitochondria-derived structures within autophagosomes. The quadrimembrane structure and small content of BNIP3 vesicles should make them lighter than Mito-L and therefore amenable to separation from both Mito-L and normal mitochondria based on differences in their respective densities. To test this, we performed iodixanol-density gradient ultracentrifugation of homogenates from drug-treated HeLa cells. As shown in Fig. 3C, when we fractionated untreated HeLa cells, we found that the mitochondrial marker TOMM20 (a MOM protein) was distributed around the center of the fractions (∼15% iodixanol; fractions #6–8). When we treated the cells with DMOG/BafA1, additional TOMM20 peaks appeared in both lighter (#2) and heavier fractions (#11–12) than in controls. BNIP3 and the Mito-L marker TFAM showed a preferential partitioning into the lighter and heavier fractions, respectively (Fig. 3C). Importantly, we found that conditions abrogating Mito-L formation (i.e., knockdown of FIP200 or BNIP3/NIX) reduced the TFAM peak in the heavier mitochondrial fractions (Figs. 3D and S4B). We conclude therefore that the heavier mitochondrial fractions contained Mito-L. We also observed a similar shift of TFAM from the normal mitochondrial fraction to the Mito-L fraction following a 24-hour treatment with DMOG, a condition that induces Mito-L formation (Fig. S4C).

Next, we found that reducing autophagy by knocking down FIP200 reduced the intensity of the TOMM20 peak in the lighter fraction. This indicates that the lighter fraction contained BNIP3 vesicles. Importantly, we found that reduced BNIP3 levels in the lighter fractions were accompanied by increased BNIP3 levels in the normal mitochondrial fractions. This observation is consistent with our previous immunostaining data, in which the distribution of the BNIP3 signal under DMOG/BafA1 treatment shifted from a focal pattern to a tubular pattern in FIP200-deficient cells. On the other hand, double knockdown of BNIP3 and NIX was associated only with a modest decrease in the TOMM20 signal in the lighter fraction, suggesting contributions from other pathways. Although the primary contributor to the TOMM20 signal in the lighter mitochondrial fraction may not be BNIP3/NIX-mediated mitophagosomes, we did observe a preferential partitioning of BNIP3 into the lighter mitochondrial fraction rather than the Mito-L fractions. We will therefore refer to the lighter mitochondrial fraction as the BNIP3 fraction.

Using this method of biochemical partitioning, we also examined the distribution of mtDNA under the untreated and DMOG/BafA1-treated conditions. As shown in Fig. 3C, mtDNA appeared in the normal mitochondrial fractions in the untreated condition. Upon DMOG/BafA1 treatment, however, the mtDNA shifted from the normal mitochondrial fractions to the Mito-L fraction but not the BNIP3 fraction. This again demonstrates the preferential partitioning of mtDNA into Mito-L over BNIP3 vesicles upon activation of the HIF1 program.

### Mito-L forming processes protect mtDNA from mitophagic degradation

Next, we tested how Mito-L contributes to protecting mtDNA from degradation during excessive mitophagy. Based on our model (Fig. S4A), long tubular mitochondria are required for Mito-L formation before mitophagy induction. Therefore, advanced mitochondrial fragmentation is expected to prevent Mito-L formation. To test this hypothesis, we fragmented mitochondria by knocking down OPA1, an essential gene for the mitochondrial fusion process (Fig. S4A, D-E). We then examined Mito-L formation in OPA1-deficient cells in the presence of DMOG. As expected, the fragmented mitochondria in these cells failed to form Mito-L efficiently (Figs. 4A, S4A and S4E).

**Fig. 4.**
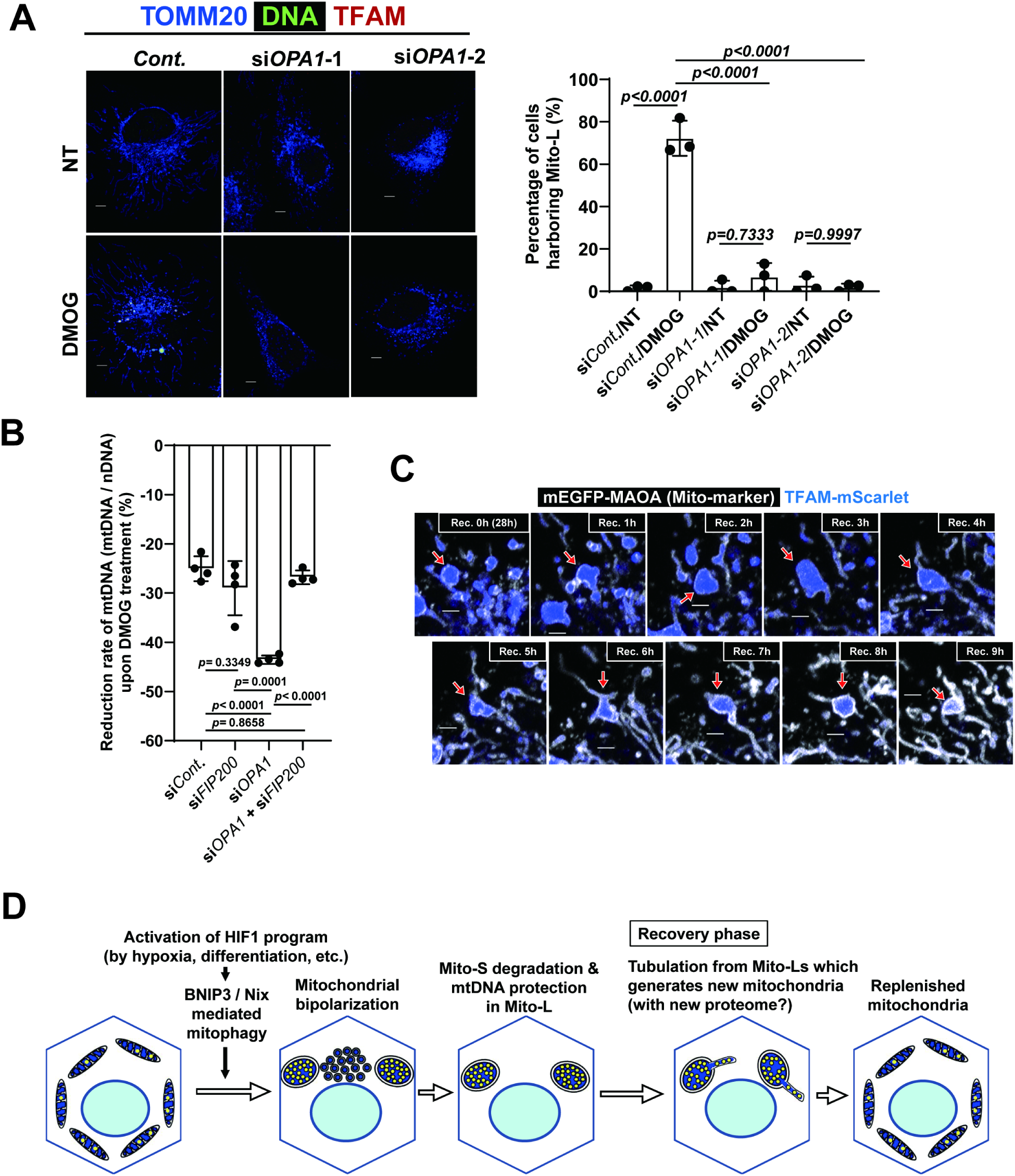
Mito-L forming processes protect mtDNA from mitophagic degradation. (**A**) HeLa cells treated with the indicated siRNAs were subjected to DMOG (1 mM) treatment for 30 h. Mito-L formation was visualized via immunostaining with the indicated antibodies. The right graph indicates the percentage of cells harboring Mito-L. (**B**) HeLa cells treated with the indicated siRNAs (siOPA1: #1, siFIP200: #1) were subjected to DMOG (1 mM) treatment for 30 h. mtDNA and nuclear DNA (nDNA) were quantified via qPCR performed on total extracted DNA. The mtDNA level was calculated relative to the nDNA level. In the graph, changes in mtDNA induced by DMOG treatment are presented as fold changes within each siRNA-treated group. (**C**) Mito-L were induced in HeLa cells transiently expressing TFAM-mScarlet and EGFP-MAOA by DMOG treatment (28 h). After removal of DMOG from the culture medium, the cells were subjected to time-lapse fluorescent imaging. (**D**) A schematic showing how Mito-L formation protects mtDNA from destruction during extensive mitophagy and how new mitochondria are formed from Mito-L during the recovery phase. In panels A and B, the data are presented as means ± SD from three (A) or four (B) independent samples. Significant differences were calculated using two-way ANOVA (A) or one-way ANOVA (B) with Tukey’s tests for multiple comparisons. Scale bars, 5 μm (A), 2 μm (C).

We next measured mtDNA levels following DMOG treatment. As shown in Fig. 4B and Fig. S4F, control siRNA-treated cells exhibited a modest reduction in mtDNA levels (- 25.1%) following DMOG treatment compared to non-treated cells. In contrast, OPA1-deficient cells treated with DMOG showed a significant reduction in mtDNA levels (- 43.5%) than untreated OPA1-deficient cells. Importantly, the additional inhibition of autophagy via FIP200 knockdown reversed this effect of OPA1 knockdown. HIF1 accumulation, BNIP3 and NIX upregulation, and TOMM20 reduction following DMOG treatment remained comparable between control siRNA-treated cells and OPA1 siRNA-treated cells. Additionally, BNIP3 and NIX accumulation increased in autophagy-repressed conditions (i.e., in the FIP200-deficient groups), as expected (Fig. S4G).

These results indicate that when Mito-L formation is defective, mtDNAs are insufficiently protected from degradation during mitophagy. Furthermore, OPA1 knockdown promoted the incorporation of mtDNA into mitophagosomes. Consistent with this, OPA1 knockdown increased the number of BNIP3-positive mitophagosomes containing mtDNA per cell compared to the control group (Figs. S5A-B). Notably, BNIP3 vesicles in OPA1-deficient cells were larger than those of control cells (Figs. S5A and C). This suggests the long tubular structure of mitochondria is necessary for generating the small BNIP3-positive mitophagosomes that exclude mtDNA. Together, we conclude that Mito-L formation protects mtDNA from degradation during HR-mediated mitophagy.

Finally, we examined the fate of Mito-L during recovery from the hypoxic response. To do this, we treated HeLa cells transiently expressing TFAM-mScarlet and EGFP-MAOA with DMOG for 28 hours to induce Mito-L formation. Afterward, we washed out the DMOG by changing the medium to normal complete DMEM (Figs. 4C and S5D, Movie S4). Then, time-lapse imaging revealed multiple narrow tubules extruding from the Mito-L, causing the size of the Mito-L to decrease over time. This strongly suggests that new tubular mitochondria were generated from Mito-L during the recovery phase.

### Conclusions & Perspective

Mitochondrial DNA encodes essential subunits of the respiratory chain complex, which is why all aerobic eukaryotes have retained mtDNA throughout their long evolutional history, beginning with the acquisition of their mitochondrial ancestors (*18, 19*). It remains unclear how this retention was achieved however, particularly in conditions characterized by excessive mitophagy. In this study, we demonstrated that mammalian cells generate large, spherical mitochondria (Mito-L) enriched with mtDNA under conditions that induce mitophagy associated with the hypoxic response. Importantly, we showed that the HIF1-mediated transcription program induces Mito-L formation, and that this process requires BNIP3/NIX-mediated mitophagy, which tends to avoid engulfment of mtDNA. We also found when we abolished Mito-L formation, mtDNA nucleoids were severely degraded upon HR activation. Thus, we conclude that cells store mtDNA in Mito-L to protect it from mitophagic degradation.

Mito-L formation begins when long tubular mitochondria shrink and becomes spherical. This process requires mitophagy via BNIP3/NIX. In the presence of DMOG, BNIP3 and NIX appear as focal structures on mitochondria that are presumably pinched off from the long tubular mitochondria by autophagy. This is consistent with a previous study indicating that the autophagy process can pinch off mitochondria even in *DNM1L* knockout cells (*20*). BNIP3 vesicles preferentially incorporate mitochondrial outer and inner membrane proteins, while exhibiting limited inclusion of matrix proteins and mtDNA.This suggests continuous mitophagy via BNIP3/NIX reduces the surface area of mitochondria relative to their volume, resulting in the generation of spherical mitochondria. TFAM and mtDNA reportedly undergo liquid-liquid phase separation (LLPS) (*16, 17*). The mtDNA in spherical mitochondria, along with TFAM, may undergo LLPS and contribute to Mito-L formation. Preferential mitochondrial membrane degradation has also been reported in budding yeast, although the process is mediated by mitochondria-derived vesicles rather than mitophagy (*21, 22*). Given that this yeast system lacks components of the mitochondrial matrix, budding yeasts may use this mechanism for partial degradation of mitochondria while avoiding mtDNA degradation. The precise mechanisms by which BNIP3/NIX-mediated mitophagy excludes mtDNA remain unclear. We assume mitophagy may create a diffusion barrier that prevents mtDNA from flowing into the mitophagosome. Alternatively, the location of mitochondria engulfed by the mitophagosome may just be sufficiently distant from that of the mtDNA. Future studies will examine these possibilities.

Previous study has shown that inhibiting mitochondrial division leads to the appearance within the mitochondrial network of bulb-like structures (Mito-bulbs) enriched with mtDNA (*23*). While Mito-bulbs and Mito-L both exhibit large, spherical shapes and mtDNA enrichment, they are distinct structures. Unlike the cristae of Mito-bulbs, which are highly developed, Mito-L feature poorly developed cristae (Figs. 1H, S1H, S5E, and Movie SX). Furthermore, Mito-L formation relies on mitophagy, whereas Mito-bulb formation does not. Despite these differences, both Mito-L and Mito-bulbs store mtDNA, suggesting they may serve a similar physiological purpose in different contexts.

HR activation and BNIP3/NIX-dependent mitophagy reportedly occur not only under pathological hypoxic conditions, such as ischemia (*24, 25*), but also during physiologic conditions such as oocyte maturation (*26, 27*), cellular differentiation (*28-34*), and metabolic perturbations like succinate or fumarate accumulation (*35, 36*). Thus, HIF1-mediated mitophagy is a frequent physiological process. This type of mitophagy, because it preserves mtDNA, is crucial for preventing mtDNA loss from cells. In addition, Toren Finkel’s group used mice expressing a mitophagy reporter to demonstrate that mitophagy occurs throughout the body (*6, 37*). While the precise pathways underlying these mitophagy events have yet to be fully elucidated, we expect that they are further linked to systems involved in the preservation of mtDNA.

In contrast, PARKIN-mediated mitophagy works with PINK1 to monitor the functional state of mitochondria and selectively degrade only those that are damaged or dysfunctional (*38, 39*). This process does not occur in the absence of mitochondrial abnormalities. BNIP3/NIX-mediated mitophagy lacks this selectivity, promoting mitochondrial degradation while also safeguarding mtDNA. As a result, it can partially degrade mitochondria without causing mtDNA loss, even in the absence of mitochondrial dysfunction. In summary, while PARKIN-mediated mitophagy is predominantly associated with mitochondrial quality control, BNIP3/NIX-mediated mitophagy is better suited for overall mitochondrial reduction or remodeling, as demonstrated in erythroid differentiation (*3-5*) and neuronal development (*32*). From this perspective, identifying conditions that induce BNIP3 or NIX expression could reveal previously unrecognized instances of mitochondrial remodeling in both physiological and pathological settings.

## Acknowledgments

Takehiro Yasukawa and Mikako Yagi provided helpful advice for quantifying mtDNA. Koji Okamoto provided critical comments that enhanced our understanding of the phenomena uncovered in this paper. Eisuke Itakura shared unpublished data that contributed to the conclusions of this study. Yusaku Nakabeppu and members of his laboratory, especially Daisuke Tsuchimoto, supported the initiation of this project at Kyushu University. The FBS Core Facility at Osaka University and the Laboratory for Technical Support in the Medical Institute of Bioregulation at Kyushu University assisted with the acquisition of confocal images. The Center for Medical Research and Education in the Graduate School of Medicine at Osaka University (Katsutoshi Niwa and Ayako Goto) provided support for the use of the ultracentrifuge (Optima MAX-XP, Beckman Coulter). pcDNA3 TFAM-mScarlet was a gift from Stephen Tait (Addgene plasmid # 129573). We used [ChatGPT-3.5 and -4o, OpenAI] during the preparation of this work to improve language and readability.

## Author contributions

Conceptualization: KY

Methodology: KY, KO, AS, RK

Investigation: KY, KO

Visualization: KY, KO, AS, RK

Funding acquisition: KY, FI

Project administration: KY

Supervision: KY

Writing – original draft: KY

Writing – review & editing: KY, FI, KO

## Data and materials availability

The data required to assess the conclusions of this manuscript are included in the main text or supplementary materials. Any unique and stable reagents developed in this study can be obtained from the lead contact upon request.

## Supplementary Materials

### Materials and Methods

#### Plasmids, antibodies, siRNAs, Cells

pcDNA3 TFAM-mScarlet was a generous gift from Stephen Tait (Addgene plasmid #129573; *11*). pEGFP-MAOA was generated by insertion of a PCR-amplified DNA segment encoding the C-terminal portion of human MAOA (490 aa-527 aa) into the EcoRI & BamHI sites of pEGFP-C1.

We obtained commercial antibodies from the following sources: anti-TOMM20 Rb IgG (Santa Cruz, #sc-11415; Proteintech, #11802-1-AP), anti-TOMM20 Mo IgG (Abcam, #ab56783), anti-DNA Mo IgM (Merck Millipore, #CBL186), anti-TFAM Mo IgG (Santa Cruz, #sc-166965), anti-TFAM Rb IgG (CST, #8076; Proteintech, 22586-1-AP), anti-LAMP1 Rb IgG (CST, #9091), anti-BNIP3 Mo IgG (Abcam, #ab10433; Proteintech, #68091-1-Ig), anti-BNIP3 Rb IgG (CST, #44060), anti-NIX/BNIP3L Rb IgG (CST, #12396), anti-TIMM23 Mo IgG (BD, # 611223), anti-ATP5A1 (Proteintech, #66037-1-Ig), anti-HSP60 (Proteintech, #66041-1-Ig), anti-LC3 Mo IgG (MBL, M152-3 for ICC), anti-LC3 Mo IgG (MBL, #186-3 for WB), anti-HIF1a Rb IgG (Gene Tex, #GTX127309), anti-FIP200 Rb IgG (CST, #12436), anti-TUBB Mo IgG (Proteintech, #66240-1-Ig), anti-OPA1 Rb IgG (CST, #80471), anti-DRP1 (Proteintech, #12957-1-AP), anti-mouse IgG FCγ-specific goat IgG Alexa Fluor 488-conjugated (Jackson ImmunoResearch, #115-545-071), anti-mouse IgM m chain-specific goat IgG Alexa Fluor 647-conjugated (Jackson ImmunoResearch, #115-605-075), anti-mouse IgG Donkey IgG Alexa Fluor 647-conjugated (Jackson ImmunoResearch, #715-605-150), anti-mouse IgG Donkey IgG Alexa Fluor 488-conjugated (Jackson ImmunoResearch, #715-545-150), anti-Rabbit IgG Goat IgG Alexa Fluor 488-conjugated (Invitrogen, #A11008), Anti-rabbit IgG Donkey IgG Alexa Fluor 594-conjugated (Jackson ImmunoResearch, #711-585-152), anti-BrdU Rat IgG-Alexa Fluor 647-conjugated (Abcam, #ab220075), anti-rabbit immunoglobulins goat immunoglobulin HRP-conjugated (Dako, #P0448), and anti-mouse IgG goat IgG HRP-conjugated (Bio-Rad, #1706516). We labeled anti-LC3 Mo IgG (MBL, M152-3 for ICC) with Alexa Fluor 647 using the Alexa Fluor 647 Antibody Labeling Kit (Invitrogen, A88068) according to the manufacturer’s instructions. This antibody was used for the simultaneous detection of mitophagosomes and mtDNA.

Pre-designed and validated Silencer Select siRNAs were obtained from Thermo Fisher (BNIP3 s2061, NIX s531927, FIP200 #1 s18994, FIP200 #2 s18995, OPA1 #1 s9850, OPA1 #2 s534148, DNM1L s19568). Negative control Silencer Select #1 siRNAs were obtained from Thermo Fisher.

HeLa cells (RCB0007) and the L23immo human immortalized fibroblast cell line were provided by RIKEN BRC through the National BioResource Project of the MEXT in Japan.

#### Cell culture analyses

HeLa cells were maintained in complete DMEM [DMEM containing 4.5 g/L glucose, 584 mg/L L-glutamine (Wako #044-29765), 1 mM sodium pyruvate (Nacalai, #06977-34), 100 U/ml penicillin-streptomycin (Nacalai) and 10% (vol/vol) FBS (PAA Laboratories GmbH, A15-080)] at 37°C in 5% CO_2_. Transfection was performed according to the manufacturer’s instructions using Lipofectamine 2000 (Invitrogen, # 11668019) for plasmids or using Lipofectamine RNAiMAX (Invitrogen, # 13778100) for siRNAs. For knockdown experiments, cells were transfected with siRNAs at a final concentration of 10 nM. During incubation, cells were split twice. 72–96 h post-transfection, the cells were treated with Dimethyloxalylglycine (DMOG; Sellek; 1 mM) and/or Bafilomycin A1 (BioViotica; 100 nM). L23immo cells were maintained in POWEREDBY10 medium (GPBioSCIENCES, #6012) or DMEM as described above at 37°C and 5% CO_2_.

Cell cultures under hypoxic conditions were adjusted to an oxygen concentration of < 0.1% using the nBIONIX-3 hypoxic culture kit (Sugiyama-Gen). The cell culture medium was pre-incubated in the hypoxic kit at an oxygen concentration of less than 0.1% for 6 hours and then replaced during hypoxic treatment. Oxygen concentrations were monitored during the experiments. Within the kit, CO_2_ was maintained at around 5%.

For labeling nascent RNAs, 5-Bromouridine (Sigma, 850187-1G) was added to pre-existing medium to a final concentration of 5 mM. After culturing for 1 h at 37°C in a 5% CO_2_ incubator, the cells were fixed in 4% PFA in PBS (Nacalai, #09154-85) at 37°C for 10 min. Then, the cells were washed three times with PBS and blocked in PBS containing 5% FBS and 0.3% Triton X-100. For detection of RNA-incorporated BrU, anti-BrdU rat monoclonal IgG-conjugated with Alexa Fluor 647 (Abcam, #ab220075) was used as a primary antibody and anti-rat IgG donkey IgG-conjugated with Alexa Fluor 647 (Abcam, #ab150155) was used as secondary antibody to further enhance the resulting signal. For co-staining of BrU and DNA, anti-DNA mouse IgM (EMD Millipore, #CBL186) and anti-mouse IgM mu chain goat IgM-conjugated with Alexa Fluor 568 (Abcam, #ab175702) were used as primary and secondary antibodies, respectively.

For labeling nascent polypeptides, the procedure used by Zorkau et al. was followed with minor modifications (*40*). In brief, cells were cultured for 1 min at 37°C in cytosolic translation-blocking DMEM. This was prepared by mixing methionine/cystine-free DMEM (Gibco, #21013) with 48 mg/L L-cystine (Nacalai, #21052-22) and L-glutamine (Wako #044-29765), 1 mM sodium pyruvate (Nacalai, #06977-34), and 50 mg/mL cycloheximide (FUJUFILM, #037-20991). Then, this medium was replaced with cytosolic translation-blocking DMEM containing 500 mM L-Homopropargylglycine (L-HPG) (Jena Bioscience, #94897) and incubated for 2 hours at 37°C in a 5% CO_2_ incubator. After the incubation, the cells were washed once with mitochondria-protective buffer (MPB; 10 mM HEPES-KOH, pH 7.5, 10 mM NaCl, 5 mM MgCl_2_, 300 mM sucrose; 37°C). To remove unincorporated L-HPG, the cell membranes were semi-permeabilized in MPB with 0.005% digitonin (37°C) for 2 min. Then, the cells were washed once with 37°C MPB and fixed in 4% paraformaldehyde in PBS (37°C) for 10 min in the dark. The fixed cells were washed twice with 3% BSA/PBS (RT) and then permeabilized in PBS containing 0.5% Triton X-100 for 15 min (RT). The cells were washed twice with 3% BSA/PBS (RT) and stained via click chemistry using the Click-iT EdU Cell Proliferation Kit for Imaging (Invitrogen, #C10339) according to the manufacturer’s instructions. This permitted labeling of the azide groups of protein-incorporated L-HPGs with Alexa Fluor 594 dye.

For visualizing the potential of the mitochondrial inner membrane, cells were incubated in DMEM with DMOG (1 mM) for 30 hours. Then, after the medium was replaced with DMEM containing 500 nM Tetramethylrhodamine, ethyl ester (TMRE; invitrogen, #T669) and 1:2000 MitoBright LT Deep Red (MBLTDR; FUJUFULM, #340-92081), the cells were incubated for 15 min at 37°C in a 5% CO_2_ incubator. The cells were then washed three times with PBS (37°C). For microscopic observations, the medium was replaced with phenol red-free DMEM (Wako, #044-32955) containing 10% FCS (37°C). To obtain the TMRE and MBLTDR signals, we used an LSM980 microscopy system with AiryScan 2.0 (Zeiss) equipped with a Tokai Hit Stage-top Incubator (STXG-WELSX-SET, Tokai Hit, Japan). Because TMRE is easily bleached during observation, fields of interest were selected by referring to the MBLTDR signal. After that, the TMRE signal was obtained without pre-observation. During observation, the culture medium was maintained at 37°C and 5% CO_2_.

#### Immunofluorescence

The indicated cells were cultured on cover glass (Matsunami Glass, Thickness No. 1) manually coated with collagen type I-C (Nitta Gelatin) in DMEM. After treatment with the indicated reagents, the cells were fixed in pre-warmed fixation solution 1 (4% PFA in PBS) at 37°C for 10 min. After washing the cells with PBS three times, most of the cells were incubated overnight in blocking solution 1 (5% FBS, 0.3% Triton X-100 in PBS) at 4°C. Primary and secondary antibodies were diluted in antibody dilution solution 1 (1% BSA, 0.3% Triton X-100 in PBS).

For detection of LC3, PFA-fixed cells were washed three times with PBS. Then, their membranes were permeabilized with 0.01% digitonin (Nacalai, #19390-04) in PBS for 10 min at room temperature. After another three washes in PBS, the cells were incubated in blocking solution 2 (3% BSA in PBS) overnight. Primary and secondary antibodies for LC3 detection were diluted in antibody dilution solution 2 (1% BSA in PBS).

The stained cells were then mounted in Slowfade Diamond Antifade Mountant with or without DAPI (Thermo Fisher Scientific, #S36964, #S36963) and shielded with transparent nail polish. Fluorescent signal detection was performed on either an LSM900 or an LSM980 microscopy system both equipped with AiryScan 2.0. Microscopic images were acquired as z-stack image slices and subsequently combined into two-dimensional images using maximum intensity projection via the ZEN 3.5 software package (ZEISS). All microscopic images were processed with ZEN 3.5.

For simultaneous detection of mitophagosomes and mtDNA, HeLa cells treated with control siRNAs or two different *OPA1*-specific siRNAs for 72 hours were treated with DMOG (1 mM) and Bafilomycin A1 (100 nM) in complete DMEM (U+) [complete DMEM with 50 mg/mL uridine (Tokyo Chemical Industry, U0020)] for 11 h 10 min. Then, after SYBR-Gold (1:10,000) was added to the existing medium, the cells were further incubated for 20 min in the CO_2_ incubator. Then, the cells were washed one time with pre-warmed (37°C) complete DMEM. The cells were further cultured in complete DMEM (U+) containing DMOG (1 mM) and Bafilomycin A1 (100 nM) for 20 min. The cells were fixed with pre-warmed PFA (37°C) followed by PBS washes as described above. Cell membranes were permeabilized for 5 min in 0.005% digitonin plus 0.05% Triton X-100 in PBS at RT. After washing with PBS, the cells were incubated in blocking solution (5% BSA / PBS) at 4°C overnight. For immunofluorescence, the blocked cells were incubated in primary antibody solution [anti-BNIP3 rabbit IgG (1:250) and 1% BSA in PBS] for 1 hour at RT. After washing the cells with PBS, they were incubated in secondary antibody solution [Anti-rabbit IgG Donkey IgG Alexa Fluor 594-conjugated (1:100) and anti-LC3 Mo IgG conjugated to Alexa Fluor 647 (see the antibody section) plus 1% BSA in PBS] for 1 hour at RT. After washing the cells with PBS, dsDNA was stained by incubation in SYBR Gold (1:10,000) diluted in PBS for 5 min at RT. This was done to enhance the mtDNA signal. After washing the cells with PBS, the cells were mounted in Slowfade Diamond Antifade Mountant without DAPI.

#### Visualization of the mitochondrial inner membrane with PKmito ORANGE-FX (PKMO-FX)

Visualization of the mitochondrial cristae with PKMO-FX (Cytoskeleton, #CY-SC054) was performed as described by Chen et al. (*41*). HeLa cells were cultured as described in the "Immunofluorescence" section. The cells were maintained in complete DMEM with or without 1 mM DMOG at 37°C in a 5% CO₂ incubator for 27.5 hours. The culture medium was then replaced with complete DMEM containing 500 nM PKMO-FX and 10 μM verapamil hydrochloride (Wako, #222-00781) with or without 1 mM DMOG and incubated for 2 hours in a 5% CO₂ incubator. Afterward, the staining medium was replaced with complete DMEM with or without 1 mM DMOG, and the cells were further incubated in the CO₂ incubator for 30 minutes. After treatment, the cells were washed once with pre-warmed PBS and then fixed in pre-warmed fixation solution 1 (4% PFA in PBS) at 37°C for 15 minutes. After washing three times with PBS, the cell membranes were permeabilized with membrane permeabilization solution (0.3% Triton X-100 in PBS) at room temperature for 5 minutes. After another three PBS washes, the cells were incubated in blocking solution 2 (5% BSA in PBS) at 4°C overnight. For the detection of TOMM20 and mtDNA, anti-TOMM20 rabbit IgG (Proteintech, #11802-1-AP) and anti-DNA mouse IgM (Merck Millipore, #CBL186) were used as primary antibodies, respectively. Alexa Fluor 488-conjugated goat anti-rabbit IgG (Invitrogen, #A11008) and Alexa Fluor 647-conjugated goat anti-mouse IgM μ chain-specific IgG (Jackson ImmunoResearch, #115-605-075) were used as secondary antibodies. Both primary and secondary antibodies were diluted in antibody dilution solution 2 (1% BSA in PBS). The stained cells were mounted using SlowFade Diamond Antifade Mountant with DAPI and sealed with transparent nail polish. Fluorescent signals were detected using an LSM900 confocal microscopy system with Airyscan 2.0. Z-stack image slices were combined into two-dimensional images using maximum intensity projections via the ZEN 3.5 software package (ZEISS). Microscopic images were processed using ZEN 3.5.

#### SDS-PAGE and western blotting

Proteins were analyzed using SDS-PAGE. Samples were loaded onto a purchased SDS-PAGE gel (Wako, SuperSep Ace, #190-15001) and subjected to electrophoresis in running buffer (25 mM Tris, 0.2 M glycine, 0.1% SDS; w/v). Using transfer buffer (250 mM Tris, 1.5 M glycine, 20% methanol; v/v), the separated proteins were transferred to a PVDF membrane (Millipore, #0000216406) that had been activated by soaking in methanol before use. The transfer was confirmed with Ponceau S solution. Subsequently, the membrane was blocked at room temperature for 1 hour in blocking solution [5% skim milk (w/v) in TBST (137 mM NaCl, 2.68 mM KCl, 25 mM Tris, 0.05% Polyoxyethylene-20 Sorbitan Monolaurate (Tween-20); w/v)].

After removing the blocking solution, the membrane was incubated with primary antibody [diluted in 1% skim milk (w/v) in TBST] for 1 hour at room temperature with shaking. The membrane was washed three times for 3 minutes each in TBST and then incubated with secondary antibody [diluted in blocking solution] for 1 hour at room temperature with shaking. The membrane was again washed three times for 3 minutes each with TBST. For signal detection, the PVDF membrane was incubated for 3 minutes in a mixture of ECL-PRIME (Cytiva, #RPN2236A) solution and B solution in a 1:1 ratio, and then the signal was detected using the FUSION SOLO.7S. EDGE chemiluminescence imaging system (Vilber-Loumat) and its associated software EvolutionCapt Edge (Vilber-Loumat).

#### Fractionation

HeLa cells (2 x 10 cm dish) were cultured for 12 hours with or without 1 mM Dimethyloxalylglycine (DMOG) and 0.1 μM Bafilomycin A1 (BafA1). After washing the cells three times with PBS, they were resuspended in 0.5 mL of PBS and transferred to a 1.5 mL tube. The cells were precipitated by centrifugation (4°C, 500 x G, 5 min), and the supernatants were removed. Subsequently, the pellets were homogenized in 260 μL of homogenization buffer [225 mM mannitol, 75 mM sucrose, 0.1 mM Ethylene glycol-bis (2-aminoethylether)-N,N,N’,N’-tetraacetic acid (EGTA), 1×cOmplete-Mini, 1 mM phenylmethylsulfonyl fluoride (PMSF), 0.01% digitonin, 30 mM Tris-HCl, pH 7.4] via 60 strokes through a 26G syringe needle. The samples were then centrifuged again (4°C, 500 x G, 5 min), and the resulting supernatants were collected as post-nuclear supernatants (PNS). Portions of these samples (40 μL) were then mixed with 40 μL of 3x sample buffer [6% sodium dodecyl sulfate (SDS; w/v), 15% glycerol (v/v), 0.6 M Tris-HCl, pH 6.8], 40 μL of ultrapure water, and 5 μL of β-mercaptoethanol to prepare the PNS for analysis. Mitochondrial subtypes were separated via iodixanol gradient centrifugation. 50% OptiPrep solution [50% (w/v) iodixanol, 42 mM sucrose, 1 mM ethylenediaminetetraacetic acid (EDTA), 20 mM HEPES-KOH, pH 7.4] was diluted in dilution buffer (1 mM EDTA, 20 mM HEPES-KOH, pH 7.4) to create the gradients. The iodixanol concentration at each step and the layering of the PNS samples atop the iodixanol gradients in polycarbonate tubes are described in each respective experimental figure. Organelles and proteins from the samples were then fractionated by ultracentrifugation [55,000 rpm (201,078 x G avg), 4°C, 1 h, TLS-55 rotor] on the OPTIMA-MAX XP-ME ultracentrifuge (Beckman Coulter). Subsequently, 100 μL of the resulting solution was collected from the top layer and then each fraction was mixed with 50 μL of 3x sample buffer and 5 μL of β-mercaptoethanol and then incubated at room temperature for at least 30 minutes. For fraction 14, when the volume was less than 100 μL, it was adjusted to 100 μL with homogenization buffer before being subjected to the same treatment.

For mtDNA quantification in each fraction, DNA was purified from the whole volume of each fraction using NucleoSpin Tissue kits (TaKaRa, #740952) according to the manufacturer’s instructions. At the final step, DNAs trapped in the DNA-binding column were eluted with the same volume of elution buffer. The amount of mtDNA in each sample was then quantified by quantitative PCR (qPCR) using LightCycler 480 SYBR Green I Master mix (Roche, #04887352001) and a CFX Connect Real-Time System (Bio-Rad, #185-5201J1). For the quantification, the same volume of DNA solution per fraction was subjected to amplification with a primer set specific to mtDNA (5’-CGATGTTGGATCAGGACATC-3’ / 5’-AAGGCGCTTTGTGAAGTAGG-3’; *42*).

#### Live-cell imaging

Cells were imaged in complete DMEM at 37°C and 5% CO_2_ using an LSM980 microscopy system with AiryScan 2.0 (Zeiss) and equipped with a Tokai Hit Stage-top Incubator (STXG-WELSX-SET, Tokai Hit, Japan). For time-lapse imaging of Mito-L formation, HeLa cells were transfected with pcDNA3_TFAM-mScarlet and pEGFP-MAOA using Lipofectamine 2000 according to the manufacturer’s instructions. Five hours post-transfection, the cells were detached via trypsin-EDTA treatment. Then, the cells were spread on collagen-coated, 35-mm, glass-bottomed dishes (MATSUNAMI, #D11130H) in complete DMEM. The plates were manually coated with collagen using Cell matrix Type I-C (Nitta Gelatin, #210707) according to the manufacturer’s instructions. Fifty-six hours after DNA transfection, each dish was placed in the Tokai Hit Stage-top incubator on the LSM980. Then, the cells were treated with DMOG (1 mM) for 24 hours. During the treatment, Z-stack images of mScarlet and EGFP fluorescence were obtained on the LSM980 microscopy system with AiryScan 2.0 using a 63x objective with immersion oil (ZEISS, #4Y00-R0DY-1007-3VF3) and a digital zoom of 1.3x. Images were acquired at 10-min intervals. After image acquisition, each z-slice image was subjected to AiryScan processing followed by maximum intensity projection using the ZEN 3.5 software package. Videos were created to show 5 images (representing 50 minutes) per second and exported in the WMF format. For visualizing Mito-L dissociation, HeLa cells transiently expressing TFAM-mScarlet and EGFP-MAOA were treated with DMOG (1 mM) for 30 hours in the stage-heater on the LSM980 to allow for Mito-L formation. Then, after washing the cells with complete DMEM (37°C) three times, the cells were incubated in complete DMEM for an additional 24 hours. During this recovery time, cellular images were obtained as described above.

#### Quantification of mtDNA and nuclear DNA (nDNA)

HeLa cells were treated with the siRNAs indicated in Fig. 4B for 96 hours. Then, they were incubated for another 30 hours with or without DMOG (final 1 mM). The culture medium was changed from complete DMEM to complete DMEM (U+) 24 hours post-transfection. After 96 hours of siRNA treatment, the cells were washed three times in PBS and transferred to a 1.5 mL tube by scraping. After precipitation of the cells by centrifugation (500 x G, 5 min, 4°C), DNA was purified from the resulting cell pellets using NucleoSpin Tissue kits (TaKaRa, #740952) according to the manufacturer’s instructions. The amount of mtDNA in the extracted DNA was quantified (20 ng for mtDNA, 10 ng for nDNA) by qPCR using LightCycler 480 SYBR Green I Master mix (Roche, #04887352001) and a CFX Connect Real-Time System (Bio-Rad, #185-5201J1) as described above. For the quantification, DNAs were subjected to amplification with primer sets specific for mtDNA (5’-CGATGTTGGATCAGGA CATC-3’ / 5’-AAGGCGCTTTGTGAAGTAGG-3’, Inatomi et al, 2022) or nDNA (SLCO2B1). Both primer sets were obtained from the “Human Mitochondrial DNA (mtDNA) Monitoring Primer Set” (TaKaRa, #7246). The amount of mtDNA in each sample was expressed relative to the amount of nuclear DNA. In each graph, nontreated values of each siRNA were set to 1 and relative changes due to DMOG treatment are shown.

#### Statistics

Quantifications were performed using the Multi Gauge ver. 3.1 (Fujifilm) or Image Lab ver. 6.0 software packages. For data generated from n ≥ 3 biological replicates, one-way ANOVA was performed using GraphPad Prism (version 8.0, La Jolla California, USA, www.graphpad.com) with Tukey’s tests for multiple comparisons. Results were deemed significant if P < 0.05.

#### Writing with generative AI-assisted technologies

The authors used [ChatGPT-3.5 and -4o, OpenAI] during the preparation of this work to improve language and readability. The used prompt is “Please correct following sentences scientifically and grammatically”. After using this service, the authors conducted a thorough review and revision of the content as required and bear full responsibility for the integrity and accuracy of the published material.

#### Post-embedding CLEM

HeLa cells were cultured in complete DMEM on 35 mm ibidi grid dishes (ibidi, #81166) that had been manually coated with collagen, as described above. The cells were treated with 1 mM DMOG for 30 hours. During the treatment, cells were stained with SYBR Gold (SYBR-Go, 1:100,000) and PKMO-FX (1:1000) in the presence of 10 μM verapamil hydrochloride. After staining, the medium was replaced with complete DMEM containing either 1 mM DMOG or no DMOG, and the cells were further incubated in a CO₂ incubator for 30 minutes. Following treatment, the cells were rinsed with rinse buffer (4.5% sucrose in Sørensen phosphate buffer [PB], pH 7.35) and fixed with 2% glutaraldehyde (ELECTRON MICROSCOPY SCIENCES, #16220-P) in PB at room temperature for 15 minutes. After three additional rinses with the rinse buffer, fluorescence imaging was performed using an LSM900 microscope equipped with Airyscan 2.

The cells were then post-fixed with 1% osmium tetroxide containing 4.5% sucrose in 0.1 M PB on ice for 1 hour, followed by two rinses with distilled water. Dehydration was carried out using a chilled ethanol gradient (50%, 70%, and 90% for 5 minutes each), followed by two changes of 99.5% ethanol and two changes of absolute ethanol, each for 10 minutes. The samples were subsequently infiltrated with EPON812 resin (Luft formulation, 4:6; TAAB, #3402) and polymerized at 65 °C for 48 hours.

Target cells were retrieved according to the method described by Ohta et al. (*43*). Briefly, the polymer layer on the bottom of the dish was removed using toluene to expose the resin surface containing the cells. A 0.5 mm × 0.5 mm block containing the target cell was trimmed, and approximately 80 serial ultrathin sections (50 nm thick) were prepared using an ultramicrotome (Leica Artos 3D). The sections were collected on 50 × 24 mm coverslips.

After complete drying, fluorescence images of the bottom 25 sections were acquired using a confocal laser scanning microscope (Leica Stellaris 5). The sections were then contrasted with 1% aqueous uranyl acetate, followed by modified Sato’s lead solution. Finally, a 2 nm-thick plasma osmium coating was applied to reduce charging during electron microscopy. The serial sections were subsequently imaged with a field emission scanning electron microscope (FE-SEM; JEOL JSM-IT800). Correlative analysis between the light and electron microscopy images was performed as needed. A three-dimensional model of Mito-L from a DMOG-treated HeLa cell (30 h) was obtained by aligning confocal laser scanning microscopy (CLSM) stacks with scanning electron microscopy (SEM) stacks. The mitochondrial inner membrane (red), cristae (blue), and mtDNA (yellow) were reconstructed through manual segmentation.

---

**Fig. S1.**
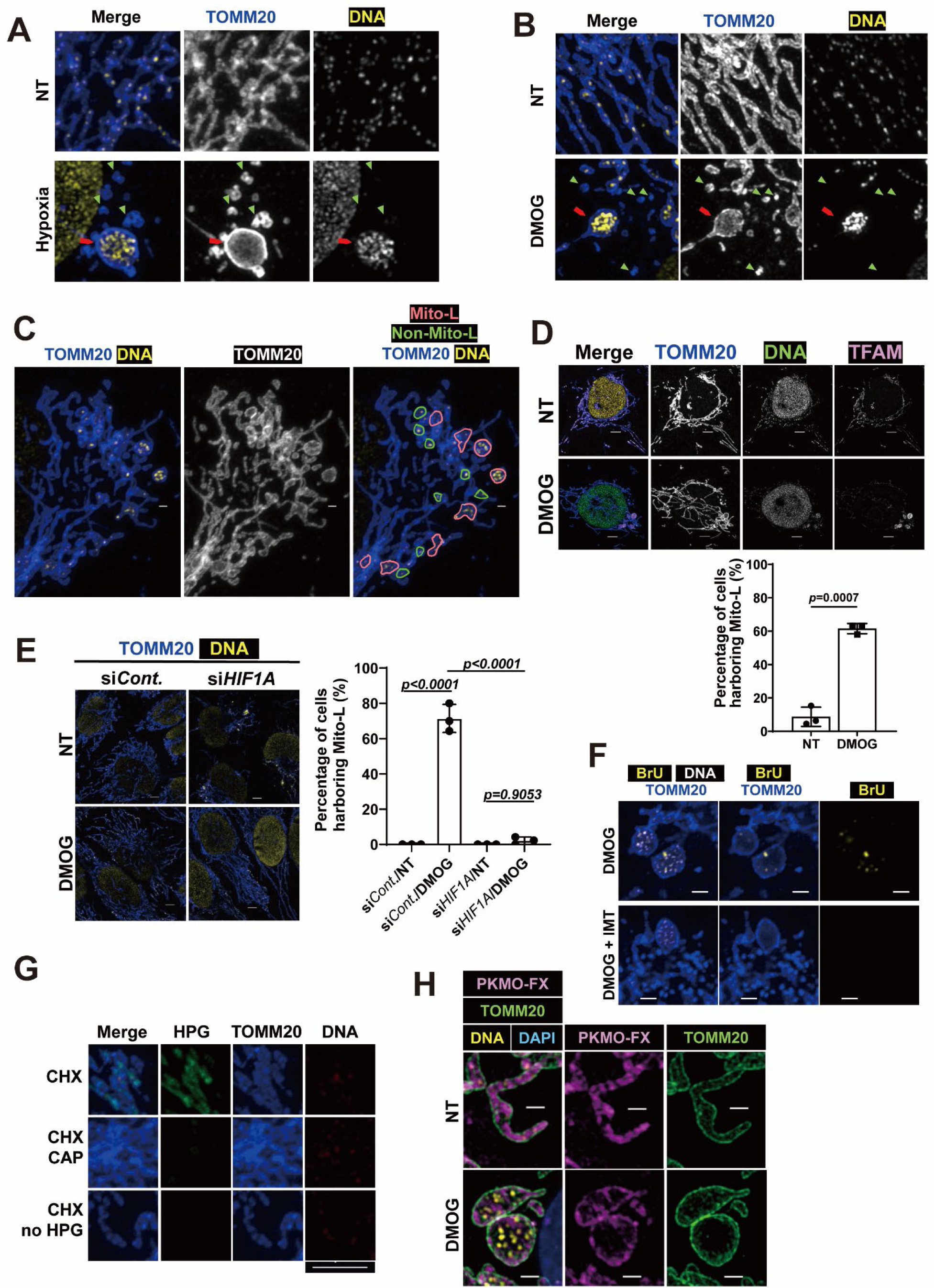
The HIF1 program facilitates Mito-L formation (supporting figure for Fig. 1). (**A, B**) High magnification images of TOMM20 and DNA staining in the same experiment as in Fig. 1A and B. Arrows and arrowheads indicate Mito-L and Mito-S, respectively. (**C**) Mito-L are defined as mitochondria swollen into a spherical shape and containing two or more mtDNA copies. Examples of Mito-L are surrounded by purple lines. Non-Mito-L structures resembling Mito-L are surrounded by green lines. (**D**) L23immo cells were subjected to DMOG (1 mM) treatment for 30 h. Mito-L formation was visualized by immunostaining with the indicated antibodies. The right graph indicates the percentage of cells harboring Mito-L. The data are presented as means ± SD from three independent samples. Statistical analysis was performed using two-tailed Student’s t-tests. P values are indicated in the graph. (**E**) HeLa cells treated with the indicated siRNAs were subjected to DMOG (1 mM) treatment for 30 h. Mito-L formation was visualized by immunostaining with the indicated antibodies. The right graph indicates the percentage of cells harboring Mito-L. The data are presented as means ± SD from three independent samples. Significant differences were calculated using two-way ANOVA with Tukey’s tests for multiple comparisons. (**F**) HeLa cells were subjected to DMOG (1 mM) treatment for 30 h. Cristae membranes were visualized with PKmito ORANGE FX (PKMO-FX), and DNA and TOMM20 were detected with the corresponding antibodies. PKMO-FX and TOMM20 background signals were subtracted. Scale bars, 1 μm (C, F), 2 μm (D), 5 μm (E).

**Fig. S2.**
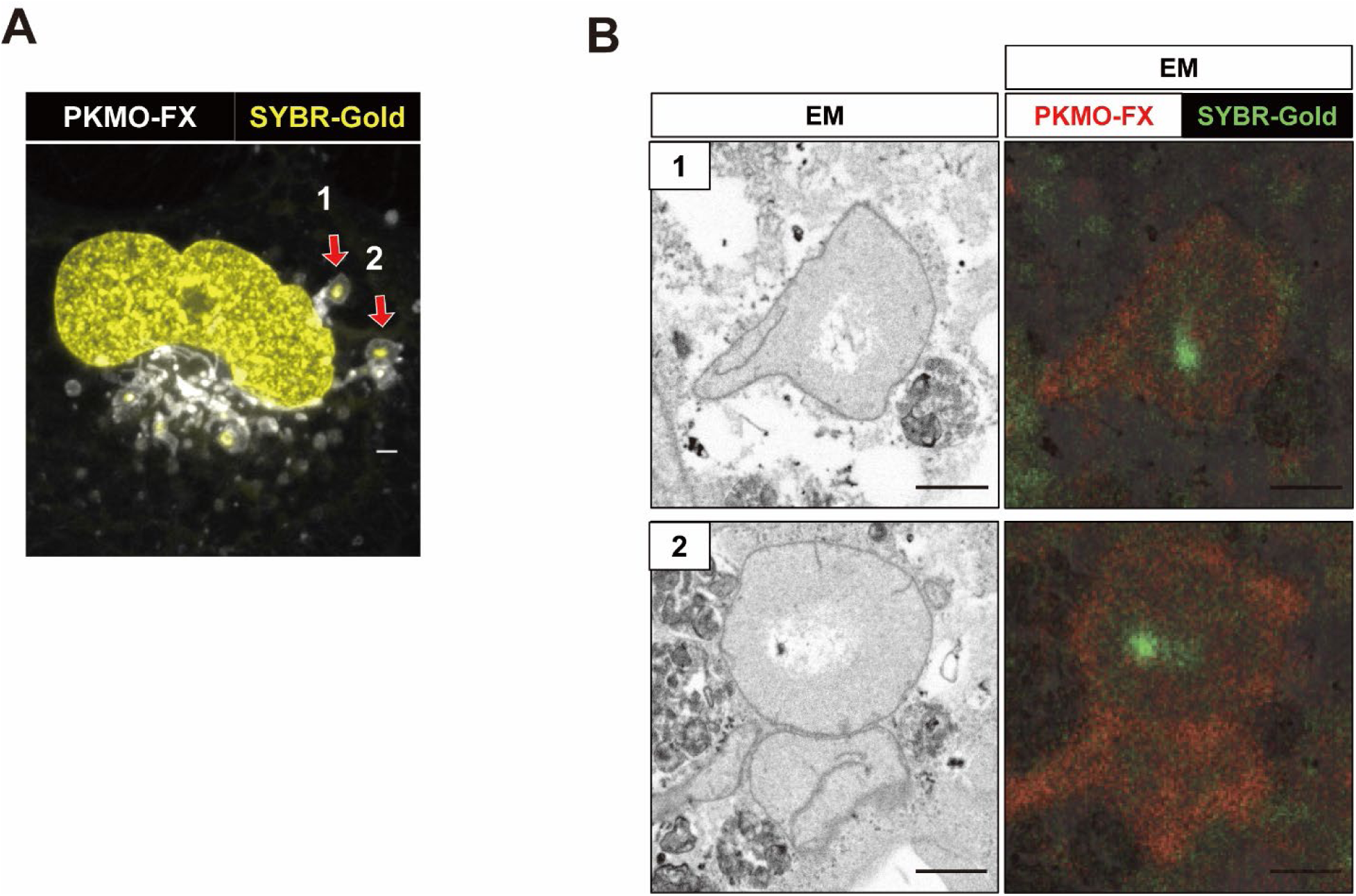
Mito-Ls exhibit poorly developed cristae (supporting figure for Fig. 1). CLEM analysis of HeLa cells treated with DMOG 30 h. (**A**) Pre-embedding confocal fluorescence image of DMOG-treated HeLa cells. DNA and the mitochondrial inner membrane were stained with SYBR Gold (green) and PKMO-FX (red), respectively. Red arrows indicate Mito-Ls analyzed in (B). (**B**) CLEM images of the Mito-Ls shown in (A). Electron micrographs (left) are merged with post-embedding confocal fluorescence images of SYBR Gold (green) and PKMO-FX (red) (right). Scale bars, 1 μm (B), 2 μm (A)

**Fig. S3.**
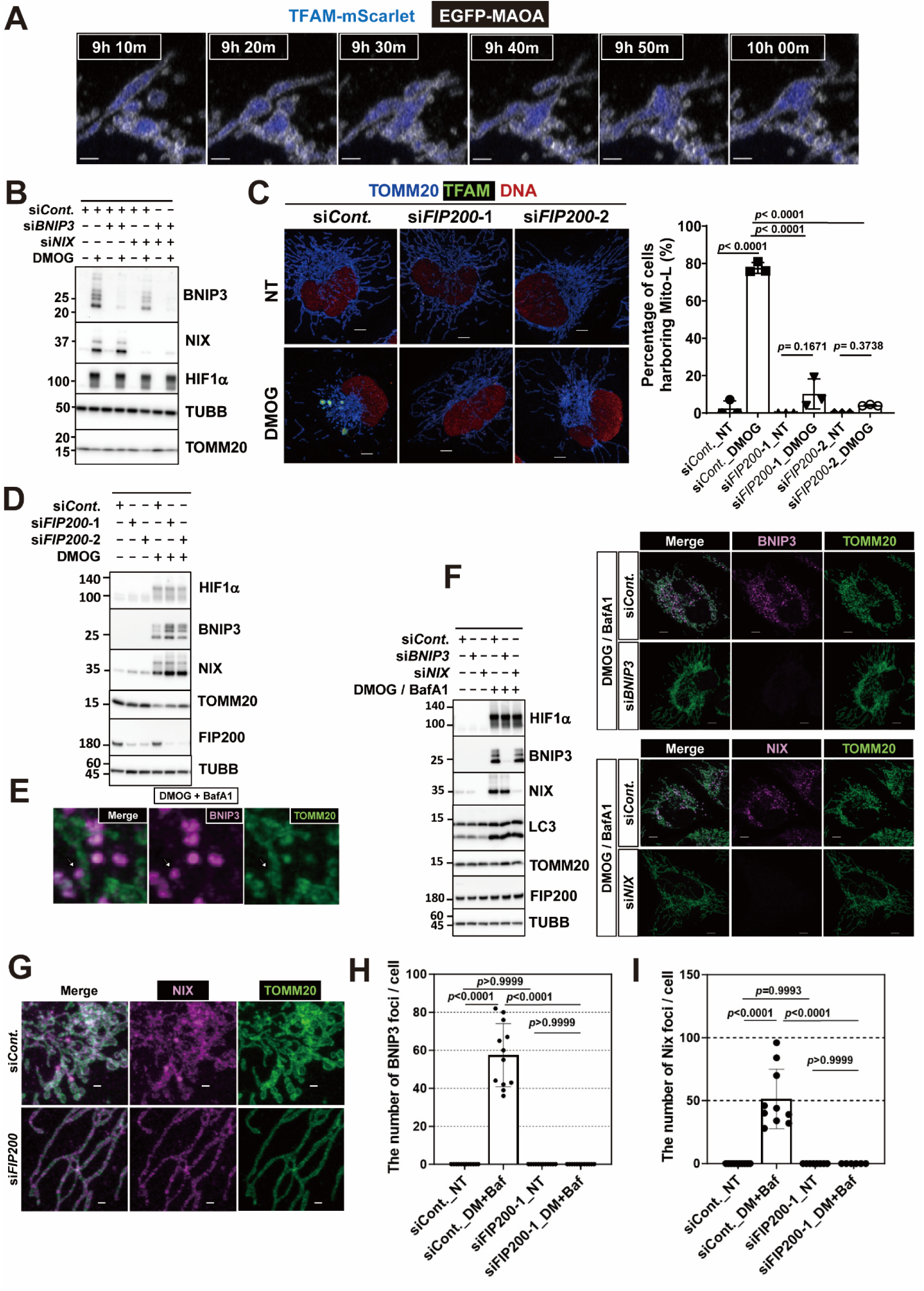
Mito-L formation depends on BNIP3/NIX-mediated mitophagy (supporting figure for Fig. 2). (**A**) Time-lapse imaging of fusion between mid-sized spherical mitochondria under DMOG treatment. These data were obtained in the same way as those in Fig. 2A. (**B**) HeLa cells treated with the indicated siRNAs were subjected to DMOG (1 mM) treatment for 30 h. Protein expression and knockout efficiency were analyzed by western blot using the indicated antibodies. (**C, D**) HeLa cells treated with the indicated siRNAs were subjected to DMOG (1 mM) treatment for 30 h. Mito-L formation and protein expression were visualized with the indicated antibodies via immunostaining (C) and western blot (D), respectively. The graph attached to the right side of (C) indicates the percentage of cells harboring Mito-L from three independent experiments. Significant differences were calculated using two-way ANOVA with Tukey’s tests for multiple comparisons. (**E**) HeLa cells were treated with DMOG (1 mM) and BafA1 (100 nM) for 12 h. Immunostaining was performed with the indicated antibodies. (**F**) HeLa cells treated with the indicated siRNAs were subjected to treatment with DMOG (1 mM) and BafA1 (100 nM) for 12 h. The specificity of the indicated antibodies was confirmed via western blot (left) and immunostaining (right). (**G**) HeLa cells treated with the indicated siRNAs were subjected to treatment with DMOG (1 mM) and BafA1 (100 nM) for 12 h. BNIP3 distribution was visualized by immunostaining with the indicated antibodies. (**H-I**) The number of BNIP (H) or NIX foci (I) per single cell under the indicated conditions as shown in Fig. 2F, S2G. The data are presented as means ± SD from different cells. Significant differences were calculated using two-way ANOVA with Tukey’s tests for multiple comparisons. The numbers of analyzed cells were as follows: [BNIP3 foci] siCont. NT (12), siCont. DM+Baf (12), siFIP200-1 NT (12), siFIP200-1 DM+Baf (12). [NIX foci] siCont. NT (15), siCont. DM+Baf (18), siFIP200-1 NT (8), siFIP200-1 DM+Baf (10). Scale bars, 1 μm (A, G), 5 μm (C, F).

**Fig. S4.**
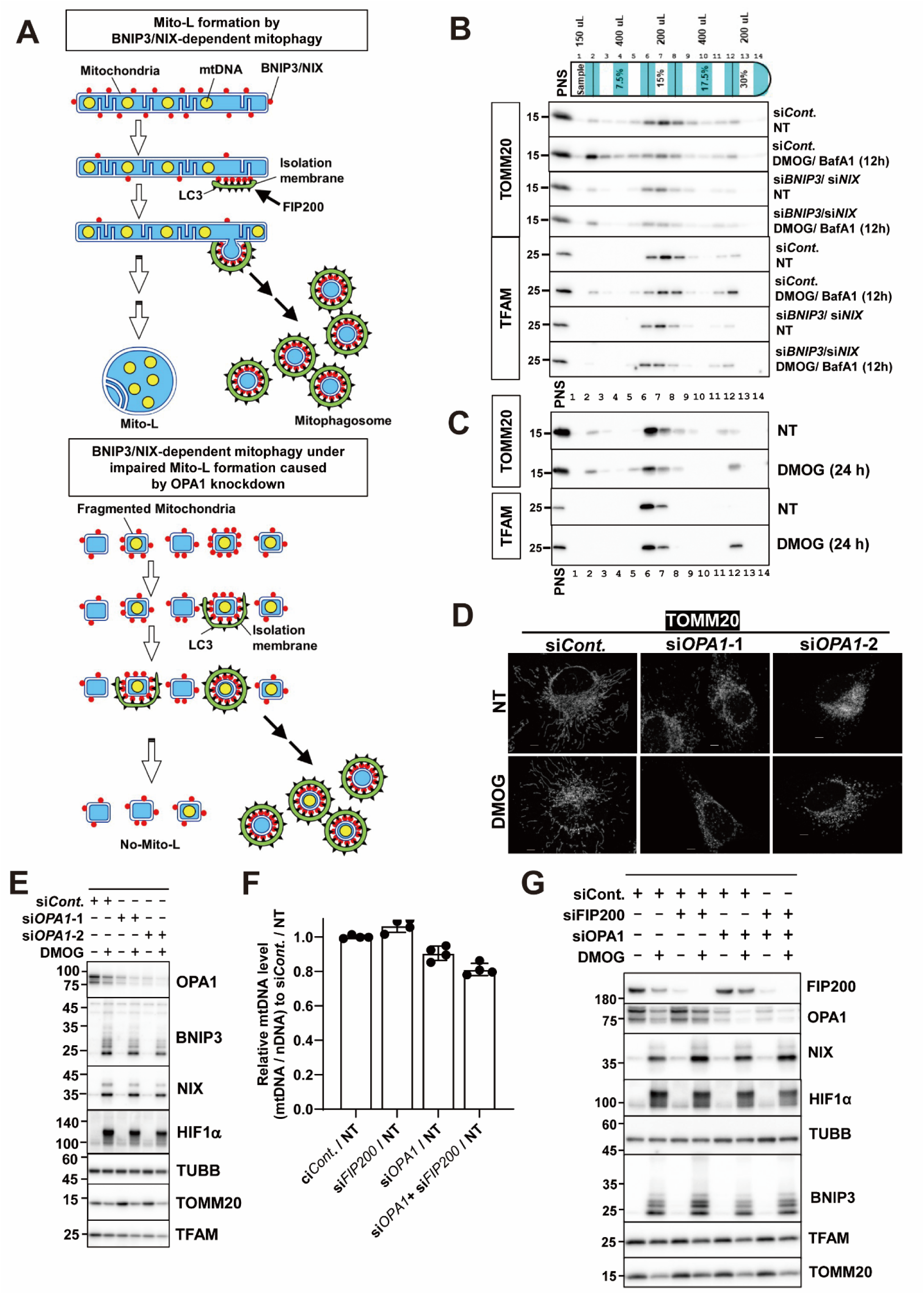
Mitophagosomes containing BNIP3 tend to omit mtDNA (supporting figure for Figs. 3 and 4). (**A**) Upper panel: Schematic model illustrating how BNIP3/NIX-dependent mitophagy contributes to Mito-L generation. Lower panel: Hypothetical model of BNIP3/NIX-dependent mitophagy under impaired Mito-L formation induced by OPA1 knockdown. (**B**) HeLa cells treated with the indicated siRNAs were subjected to treatment with DMOG (1 mM) and BafA1 (100 nM) for 12 h. Cell homogenates were separated and analyzed as in Fig. 3C. (**C**) HeLa cells were subjected to treatment with DMOG (1 mM) for 24 h. Cell homogenates were separated and analyzed as in Fig. 3C. (**D, E**) HeLa cells treated with the indicated siRNAs were subjected to treatment with DMOG (1 mM) for 30 h. The specificity of the indicated antibodies was confirmed by immunostaining (D) and western blot (E). (**F**) Relative mtDNA levels (mtDNA/nDNA) in non-treated cells as in Fig. 4B. These values are indicated relative to the mtDNA level of cells treated with control siRNAs. The experimental conditions match those of Fig. 4C. (**G**) HeLa cells treated with the indicated siRNAs were subjected to treatment with DMOG (1 mM) for 30 h. Protein expression and knockout efficiency were analyzed by western blot using the indicated antibodies. Scale bars, 5 μm (D).

**Fig. S5.**
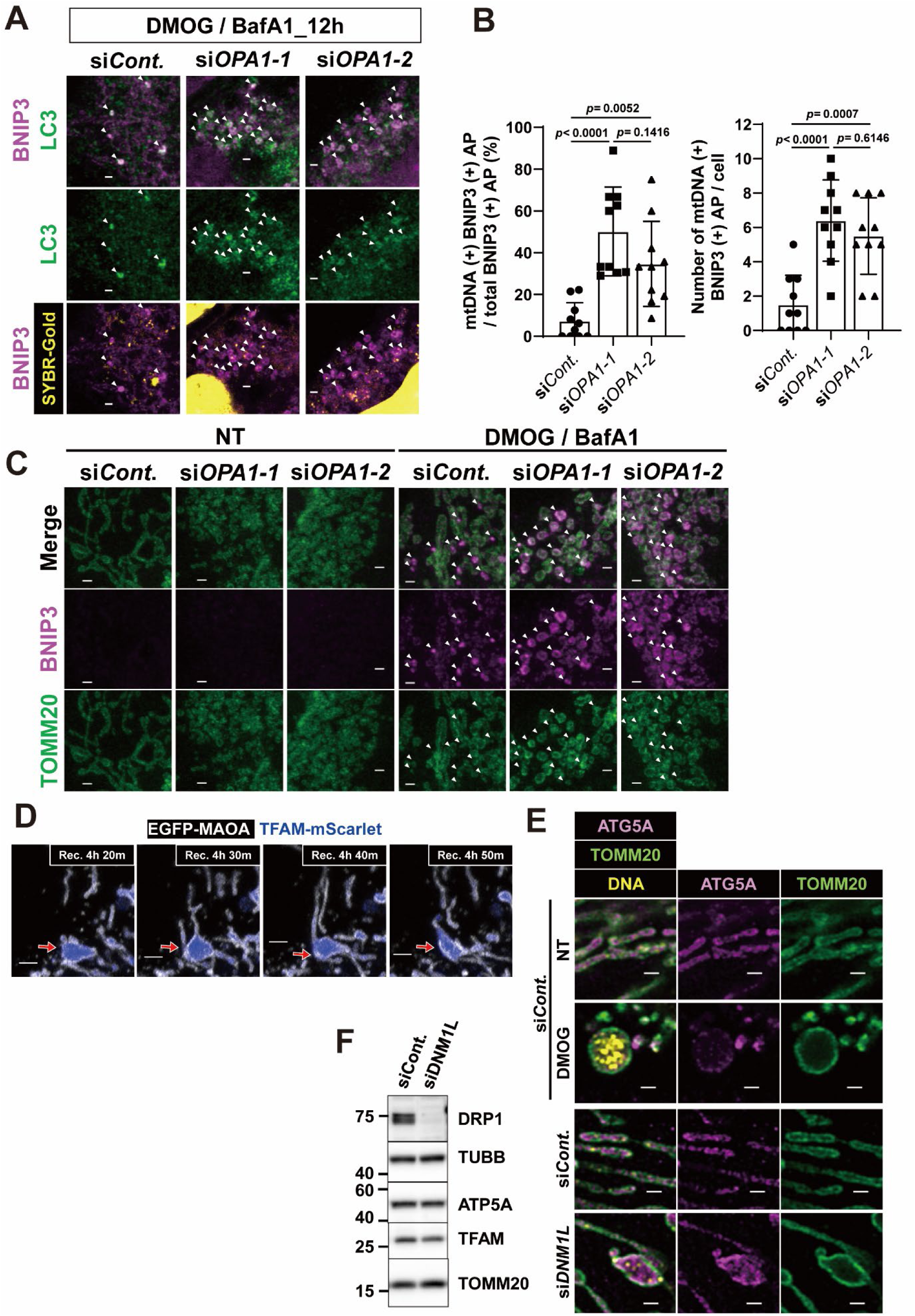
Mito-L protects mtDNA from mitophagic degradation (supporting figure for Fig. 4). (**A**) HeLa cells treated with or without DMOG (1 mM) and BafA1 (100 nM) for 12 hr were additionally treated with the indicated siRNAs. Then, the number of mitophagosomes containing BNIP3 was compared. BNIP3 and LC3 were detected using the corresponding antibodies, and dsDNA was visualized with SYBR-Gold. The arrowheads indicate BNIP3(+)/LC3(+) structures [BNIP3(+) AP]. (**B**) The left graph indicates the percentage of mtDNA(+) BNIP(+) AP relative to the total number of BNIP(+) AP in a single cell. The right graph indicates the number of mtDNA(+) BNIP(+) AP per cell. These data were collected from 10 cells per condition in the experiment shown in Fig. S4A. All cell images were obtained as a single confocal plane. The data are presented as means ± SD from different cells. Significant differences were calculated using one-way ANOVA with Tukey’s tests for multiple comparisons. (**C**) Comparison of BNIP3-enclosing structures in the same set of experiments as shown in Fig. S4A. Immunostaining was performed with the indicated antibodies. Arrowheads indicate BNIP3-positive mitochondria. (**D**) The other time-lapse images show the regeneration of narrow tubular mitochondria from Mito-L during recovery from excessive mitophagy. (**E, F**) HeLa cells treated with the indicated siRNAs. For inducing Mito-Ls, control siRNA-treated cells were treated with DMOG (1 mM) for 30h. For inducing Mito-bulbs, the cells were treated siRNA for DNM1L. Immunostaining (E) and western blotting (F) were performed with the indicated antibodies. Cristae were visualized with anti-ATP5A antibody in (E). Scale bars, 1 μm (A, B, E), 2 μm (D).

**Movie. S1. Three-dimensional model of Mito-L.** A representative three-dimensional model of Mito-L from a DMOG-treated HeLa cell (30 h) is shown. Image stacks acquired using confocal laser scanning microscopy (CLSM) and scanning electron microscopy (SEM) were aligned for reconstruction. The mitochondrial inner membrane (red), cristae (blue), and mtDNA (yellow) were reconstructed by manual segmentation. The scale bar ranges from 10 μm to 2 μm.

**Movie. S2. Time-lapse imaging of Mito-L formation upon DMOG treatment.** TFAM-mScarlet and EGFP-MAOA were transiently expressed in HeLa cells to visualize mitochondrial nucleoids and MOM, respectively. After the transfection, the cells were treated with DMOG (1 mM). During DMOG treatment, Z-stack images of mScarlet fluorescence and EGFP fluorescence were obtained at 10-minute intervals. After acquisition of the images, each z-slice image was subjected to AiryScan processing and maximum intensity projection using the ZEN 3.5 software package. Videos were created to show 5 images (representing 50 minutes) per second.

**Movie. S3. Time-lapse imaging of mitochondrial fusion events as Mito-L form under DMOG treatment.** TFAM-mScarlet and EGFP-MAOA were transiently expressed in HeLa cells to visualize mitochondrial nucleoids and MOM, respectively. After the transfection, the cells were treated with DMOG (1 mM). During the DMOG treatment, Z-stack images of mScarlet fluorescence and EGFP fluorescence were obtained at 10-minute intervals. After acquisition of the images, each z-slice image was subjected to AiryScan processing and maximum intensity projection using the ZEN 3.5 software package. Videos were created to show 5 images (representing 50 minutes) per second.

**Movie. S4. Time-lapse imaging of Mito-L dissociation after DMOG removal.** TFAM-mScarlet and EGFP-MAOA were transiently expressed in HeLa cells to visualize mitochondrial nucleoids and MOM, respectively. After the transfection, the cells were treated with DMOG (1 mM) for 30 hours to allow for Mito-L formation. After washing, the cells were incubated in complete DMEM for an additional 24 hours. During this recovery time, cellular images were obtained and processed as with Movie S2.

